# Integration of constraint-based modelling with faecal metabolomics reveals large deleterious effects of *Fusobacteria* species on community butyrate production

**DOI:** 10.1101/2020.09.09.290494

**Authors:** Johannes Hertel, Almut Heinken, Filippo Martinelli, Ines Thiele

## Abstract

Integrating constraint-based community modelling with population statistics, we introduce new theoretical concepts for interrogating the metabolic functions of the microbiome, applying them to a public metagenomic dataset consisting of 365 colorectal cancer cases (CRC) and 251 healthy controls. We found that 1) glutarate production capability was significantly enriched in CRC microbiomes and mechanistically linked to lysine fermentation in *Fusobacteria* species, 2) acetate and butyrate production potentials were lowered in CRC, 3) Fusobacteria presence had large negative ecological effects on community butyrate production in CRC and healthy controls. Validating the model predictions against faecal metabolomics, our *in silico* frameworks correctly predicted *in vivo* species metabolite correlations with high accuracy. In conclusion, highlighting the value of combining statistical association studies with *in silico* modelling, this study delivers insights on the metabolic role of *Fusobacteria* in the gut, while providing a proof of concept for the validity of constraint-based community modelling.

## Introduction

The gut microbiome with its trillions of bacteria contributes crucially to human metabolism in health and disease (Clemente et al., 2012). It generates otherwise inaccessible nutrients (Shafquat et al., 2014), inactivates and activates drugs (Wilson and Nicholson, 2017), and produces potentially harmful metabolites (Yuan et al., 2019). Recent advances in sequencing techniques have given rise to a wealth of insights into patterns of gut microbiome composition, revealing that the gut microbiome is a correlate of many human diseases (Lynch and Pedersen, 2016). Besides results stemming from observational human cohort studies, an impressive number of experimental studies on animal models have resulted in insight into the mechanisms by which the gut microbiome interacts with the host organism (Douglas, 2019). Specifically, bacterial fermentation pathways play a key role in mediating host-microbe metabolic interactions. Short chain fatty acids (SFCAs), namely acetate, butyrate, and propionate, arise from gut microbial fermentation of dietary fibre (Koh et al., 2016). Microbial fermentation of protein also results in short chain fatty acid production but mostly result in branched-chain fatty acids, such as isobutyrate, 2-methylbutyrate, and isovalerate (Smith and Macfarlane, 1997). SFCAs, especially butyrate, directly modulate host physiology by serving as signalling molecules (Koh et al., 2016). For instance, they act as histone deacetylase (HDAC) inhibitors and bind to G protein-coupled receptors (GPCRs) (Johnstone, 2002).

Increasing evidence points towards the gut microbiome contributing to colorectal cancer (CRC) through its metabolome, in particular through alterations in SFCA metabolism (Louis et al., 2014; Tilg et al., 2018). Butyrate is protective against CRC since it is both a potent anti-tumour and anti-inflammatory agent (Chang et al., 2014) mediated by its HDAC-inhibiting effects (Flint et al., 2012). Moreover, butyrate serves as the main carbon source for healthy colonocytes but not for tumour cells (Koh et al., 2016). Consistently, multiple studies have reported a decrease in butyrate-producing bacteria in CRC patients (Koh et al., 2016). On the other hand, the gut microbiome produces potentially genotoxic metabolites, such as hydrogen sulfide and secondary bile acids (Niederreiter et al., 2018), contributing potentially to CRC pathogenesis. A number of species have been implicated in the pathogenesis of CRC, such as *Fusobacterium nucleatum, Escherichia coli, Bacteroides fragilis, Gemella morbillorum, Parvimonas micra*, and *Solobacterium moreii* (Tilg et al., 2018). Moreover, a microbial signature encompassing 29 species was predictive for CRC (Wirbel et al., 2019). Hence, it has been suggested that the gut microbiome could serve as a prognostic and diagnostic marker (Thomas et al., 2019; Wirbel et al., 2019). Additionally, the microbiome changes in its composition during the progression of the disease (Yachida et al., 2019), while playing an important role in promoting resistance to chemotherapy (Yu et al., 2017) or modifying them in a toxic way (Alexander et al., 2017).

The faecal metabolome is considered to be a readout of the functional capabilities of the gut microbiome (Yachida et al., 2019; Zierer et al., 2018). Consequently, changes in faecal metabolome profiles in CRC have also been linked to altered microbial abundance patterns via statistical association studies (Kim et al., 2020; Koeth et al., 2013; Xu et al., 2020). Yet, it remains challenging to identify the mechanisms by which the microbiome changes the metabolome, as statistical associations may be caused by indirect effects and confounding (Noecker et al., 2019; Shaffer et al., 2017). Moreover, as species share metabolic capabilities and functions even across different phyla (Magnusdottir et al., 2017), it is by no means clear that a change in composition will result in a change in metabolic functions. In consequence, two gut microbial communities may look drastically different regarding their species composition, while they may be largely equivalent in terms of metabolic functions, complicating interpretations of metagenomics studies. As the gut microbiome acts as a complex ecosystem where species have to be understood in their role within communities, systems biology approaches seem to be best suited to tackle the problem of translating patterns of species abundance into patterns of metabolic function (Noecker et al., 2019).

Herein, we applied constraint-based reconstruction and analysis (COBRA) to map species abundance patterns onto patterns of metabolic functions (Heirendt et al., 2019). COBRA represents a scalable systems biology computational modelling approach, widely applied in the field of microbiome research (Chng et al., 2020; Garza et al., 2020; Henson et al., 2019; Thiele et al., 2020). Its strengths of integrating genomic data with condition specific constraints are specifically designed to deliver on the task of characterising metabolic functions of microbial communities (Orth et al., 2010). Accordingly, we utilised metabolic reconstructions of hundreds of gut microbes (Magnusdottir et al., 2017) in combination with community modelling (Baldini et al., 2018) to predict metabolic outputs of microbial communities as demonstrated previously (Heinken et al., 2019). Based on a recently published metagenomics data set of a colon cancer case-control study (Yachida et al., 2019), we successfully validate then our predictions via integrating them with faecal metabolomic measurements from the same study. Crucially, we demonstrate that AGORA-based community modelling can correctly predict the empirical species-metabolite association patterns for butyrate and glutarate. Thereby, we demonstrate the validity of COBRA community modelling in a proof-of-principle analysis, providing novel insights into the role of Fusobacteria in CRC.

## Results

To translate microbiome abundance pattern into patterns of metabolic functions, we applied community modelling to the colorectal cancer (CRC) case-control cohort (Yachida et al., 2019), which included 616 individuals (365 CRC cases and 251 healthy controls), with metagenomic data. For each individual, a personalised microbiome model was built, appropriately contextualised with a simulated Average Japanese diet, and subsequently interrogated through flux balance analysis simulations (Methods). The simulations resulted in one model producing nothing, indicating an infeasible model specification. This case was excluded from analyses. Table 1 shows the descriptive statistics for the included cases regarding the meta-data. Table 2 displays a summary of the important theoretical concepts applied in the following analyses. The resulting personalised flux profiles were then analysed in the context of clinical parameters and metabolomic findings through population statistics modelling. Thus, this study utilizes three distinct levels of modelling (Fig. 1A): 1) The strain-specific AGORA genome-scale metabolic reconstructions, 2) the personalised COBRA community models integrating diet data and the individual’s metagenomic data resulting in individual flux profiles, and 3) the statistical modelling of populations of community models. Note that the first two steps are deterministic, while the third step is stochastic.

**Table 1:** Sample characteristics of the study.

|  | **CRC Patients**<br>**(n=364)** | **Healthy controls**<br>**(n=251)** | **p-value** |
| --- | --- | --- | --- |
| Age, mean (SD) | 62.4(9.91) | 60.81(12.64) | 0.095<sup>a</sup> |
| BMI, mean (SD) | 22.95 (3.57) | 22.67(3.04) | 0.294<sup>a</sup> |
| Female, % | 39.29% | 45.82% | 0.115<sup>b</sup> |
| Stage of the disease | HS, 10.99%<br>MP, 18.41%<br>Stage 0, 19.78%<br>Stage I/II, 30.49%<br>Stage III/IV, 20.33% | NA | NA |
| Species Richness, mean (SD) | 69.74 (18.33) | 63.91 (15.96) | <0.001<sup>a</sup> |
| # Metabolites produced, mean (SD) | 157.06 (6.67) | 156.17(7.20) | 0.123<sup>a</sup> |
| # Reactions in community models, mean (SD) | 2896.80 (99.58) | 2885.28 (105.29) | 0.190<sup>a</sup> |

CRC=Colorectal cancer, SD=Standard Deviation, HS=Healthy after surgery, MP=Multiple polyps, <sup>a</sup>p-value from Welch t-tests, <sup>b</sup>p-value from Fisher's exact test

**Table 2:** Theoretical concepts used in this study.

| Theoretical Concept | Description<sup>a</sup> | Type of modelling |
| --- | --- | --- |
| **Net metabolite production capability** | The possibility to produce a metabolite | Deterministic |
| **Net metabolite production capacity** | The amount of a metabolite (mmol/d), which can be maximally produced | Deterministic |
| **Direct species production effect** | The average contribution of a species to the net metabolite production capacity of a community | Statistical |
| **Ecological species effect** | The difference between average net metabolite production capacities of communities with a species and communities without a species after discounting the direct species net production capacity | Statistical |
| **Metabolic Equivalence** | Equivalence of two communities in terms of net metabolite production capabilities | Deterministic |
| **Metabolically Sufficient** | A species/reaction is called sufficient for a metabolite, if presence of the species/reaction within a community means that the metabolite can be produced. | Deterministic |
| **Strictly metabolically sufficient** | A species/reaction is called strictly sufficient if it is sufficient given all other sufficient species/reactions. | Deterministic |
| **Metabolically necessary** | A species/reaction is called necessary for a metabolite, if absence of the species/reaction within a community means that the metabolite cannot be produced | Deterministic |
| **Strictly metabolically necessary** | A species/reaction is called strictly necessary, if it is necessary given all other necessary species/reactions. | Deterministic |

<sup>a</sup> Formal definitions can be found in the Methods section. All definitions are conditional on the applied diet constraints.

**Figure 1:**
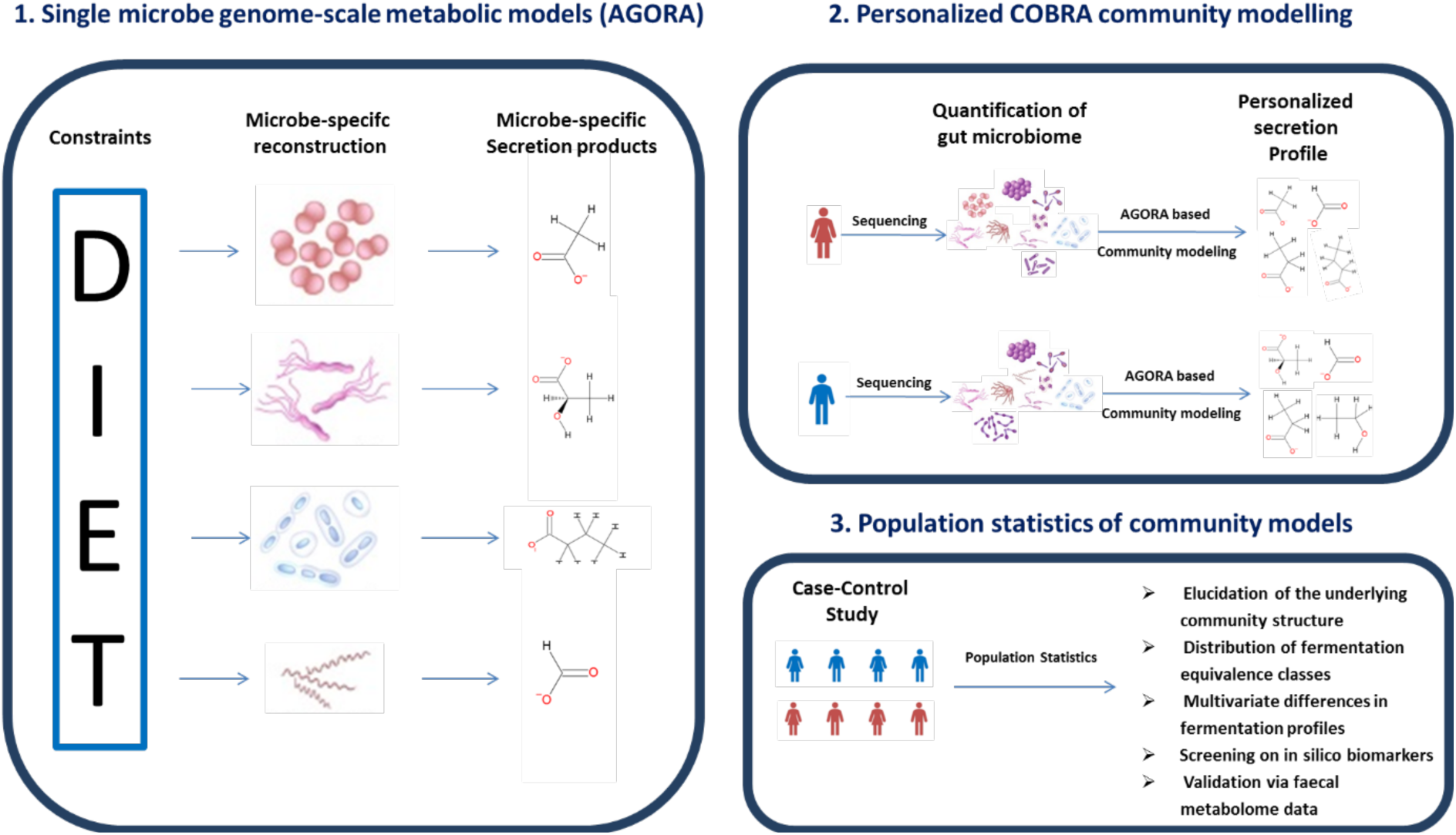
Overview over the three levels of AGORA-based community modelling used in this study.

### Microbial communities are unique in their metabolic capabilities in healthy controls and CRC cases

To gain insight into the distribution of gut microbial metabolic capabilities across samples, we explored the distribution of secretion patterns in CRC cases and controls via the concept of metabolic equivalence (see Methods). We call two communities equivalent regarding a certain set of metabolites, if the subset of metabolites with net production capacity greater zero conditional on a common diet is the same for both communities. In the AGORA resource (Magnusdottir et al., 2017), the net production capacity calculation of all 413 metabolites that are associated with exchange reactions (Noronha et al., 2019) is possible, resulting in the theoretical number of 2^413^ different equivalence classes for the whole set of metabolites. However, from these 413 metabolites, 224 metabolites were produced by no model and 90 metabolites by all models, meaning that secretion capability of 99 metabolites showed variance across the microbiome community models with 43 metabolites being produced by at least 5% of the models and maximally 95% of the models (Table S1). Despite this high level of overlapping metabolic capabilities between microbiome models, we detected 607 different equivalence classes in 615 simulated communities. Hence, microbial communities are mostly metabolically unique in their profiles of metabolic capabilities, contributing thereby to the individuality of human metabolism in health and disease.

### Glutarate production capability is enriched in CRC cases and a metabolic function unique to *Fusobacteria* sp

Next, we fitted logistic regressions to investigate whether individual metabolite secretion capabilities are enriched in CRC microbiomes controlling for age, sex, and body mass index (BMI) (Table S1 for full results). After correction for multiple testing, only the glutarate secretion capability remained significant, being clearly enriched in CRC cases (Odds ratio (OR)=2.51, 95%-confidence interval (CI)=(1.80;3.51), p=6.45e-08, FDR<0.05) (Fig. 2A). Importantly, the capability to secrete glutarate was associated with the stage of disease (p=0.003, Fig. 2B), indicating that glutarate secretion potential may be an *in silico* biomarker for CRC progression, although this result was not significant after correcting for multiple testing (FDR=0.13). Testing the association to basic covariates, we found that glutarate production capability was enriched in men (OR=1.64, 95%-CI:(1.17;2.29), p=0.004) (Fig. 2A), but not associated with age and BMI. To link the change in metabolic functions back to patterns of species abundance, we applied the concepts of necessity and sufficiency (see Methods). We identified 59 species fulfilling the criteria of being sufficient, meaning that all communities containing at least one of these species were able to secrete glutarate. From these 59 species, only seven species were strictly sufficient. Strikingly, all strictly sufficient species belonged to the genus *Fusobacterium*. Importantly, from the seven *Fusobacterium* sp., two were significantly more often detected in CRC cases (Fig. 2C). Together, these seven species were also necessary, meaning that at least one of the seven detected *Fusobacterium* species had to be present in the community for net glutarate production capacity. Hence, a community had a positive net production capacity for glutarate, if and only if *Fusobacterium* species were present.

**Figure 2:**
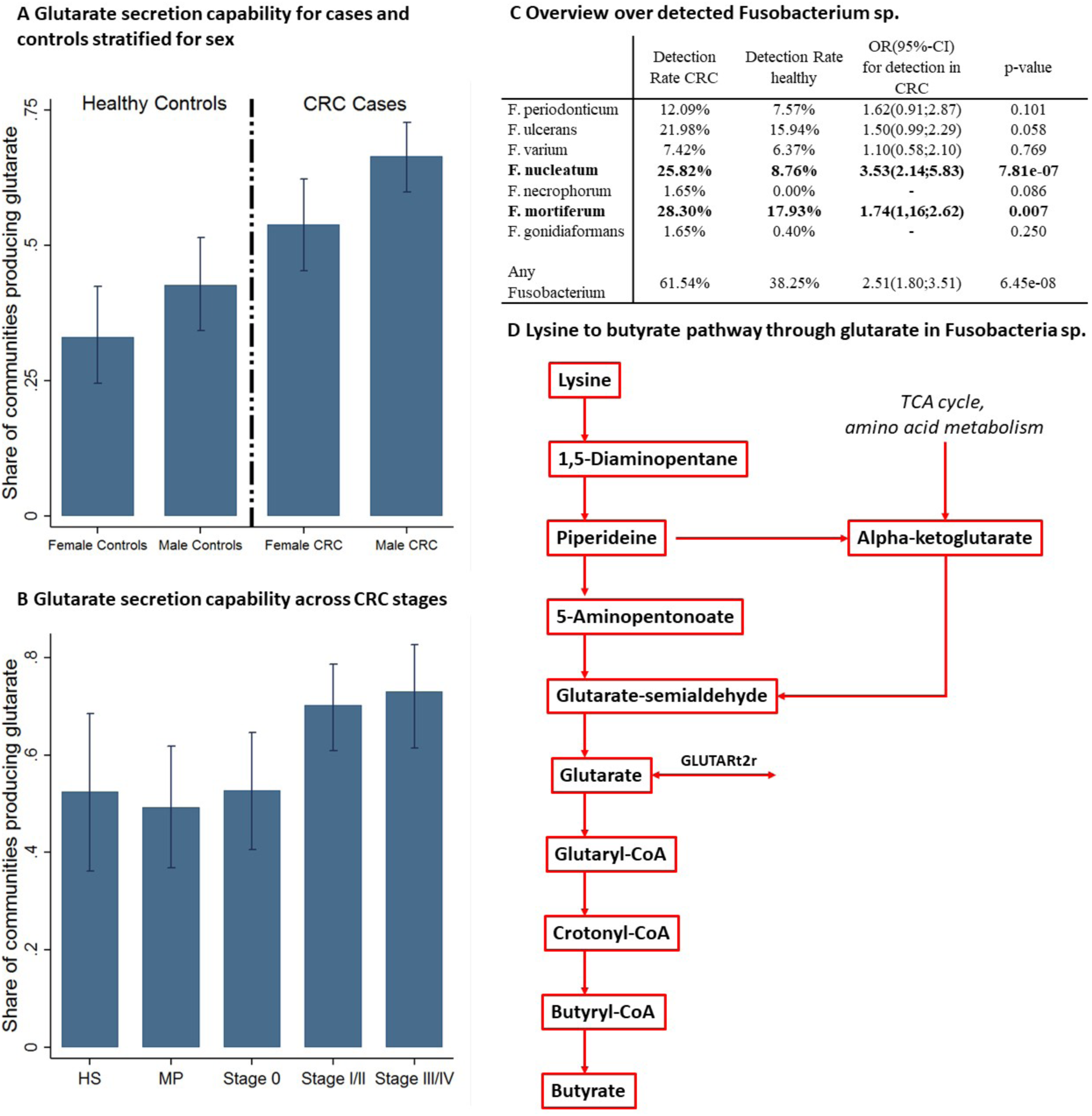
Glutarate secretion capability enrichment in CRC. **A** Bar plots with 95%-confidence intervals for the share of microbiome models with the capability to produce glutarate across the sexes and cases and controls. **B** Bar plots with 95%-confidence intervals for the share of microbiome models with the capability to produce glutarate across different stages of colorectal cancer. Late stage colorectal cancer had significantly higher shares of microbiomes with the capability to produce glutarate. **C** Statistics for the detected *Fusobacterium* species. P-values are from logistic regression adjusted for age, sex and BMI except for *F. necrophorum* and *F. gonidiaformans* where p-values were calculated from Fisher’s exact tests due to small case numbers. **D** Lysine to butyrate pathway through glutarate in *Fusobacterium* species. Note that only *Fusobacterium* species had the complete pathway including the exchange reaction for glutarate. CRC=colorectal cancer, MP=multiple polyps, HS=healthy after surgery, GLUTARt2r=Glutarate transport via proton symport, reversible.

Next, we aimed at identifying specific network properties of *Fusobacterium* species allowing for net glutarate production capabilities. Using the AGORA resource, we found that *Fusobacteria* are the only species having the complete pathway from lysine to glutarate *and* an exchange reaction for glutarate (Fig. 2D). Noteworthy, the pathway for glutarate production from lysine co-occurs with the pathway for butyrate production from glutarate (Vital et al., 2014) (Fig. 2D). Consequently, CRC microbiomes were enriched for the lysine to butyrate fermentation pathway through glutarate. In conclusion, while *Fusobacteria* sp., especially *F. nucleatum*, have been repeatedly linked to CRC, we identified a metabolic capability unique to Fusobacteria species.

### CRC microbiomes show lowered short chain fatty acid production capacities mediated by Fusobacteria presence

As glutarate is an upstream metabolite of acetate and butyrate (Buckel, 2001; Vital et al., 2014), we calculated the net secretion potential for short chain fatty acids, including propionate, by the community modelling and tested for differences in community secretion potentials between CRC cases and healthy controls. Strikingly, acetate (regression coefficient b=2.88, 95%-CI:(0.05;5.71), p=0.046) and butyrate (b=8.98, 95%-CI:(0.87;17.10), p=0.030) production potential but not propionate production potential (b=-3.61, 95%-CI:(−13.16;5.94), p=0.458), were higher in healthy controls (Fig. 3A). Noteworthy, microbiomes with Fusobacteria had lower butyrate production potential (b=-23.71, 95%-CI:(−31.52;-15.89), p=4.43e-09) in cases as well as in controls (Fig. 3B). No effect of *Fusobacteria* presence on acetate production capacities could be identified, while proprionate production potentials were higher in microbiomes with *Fusobacteria* (Fig. S1). Importantly, *Fusobacteria* presence statistically mediated the effect of CRC on butyrate production potential (Sobel-Goodman Test: Indirect effect b=5.29, 95%-CI:(2.77;7.81), p=3.79-05). Thus, our analyses provide evidence that the presence of *Fusobacteria* may be deleterious for community butyrate production potential, leading to CRC microbiomes, which are enriched for *Fusobacteria* sp., having reduced butyrate production potentials.

**Figure 3:**
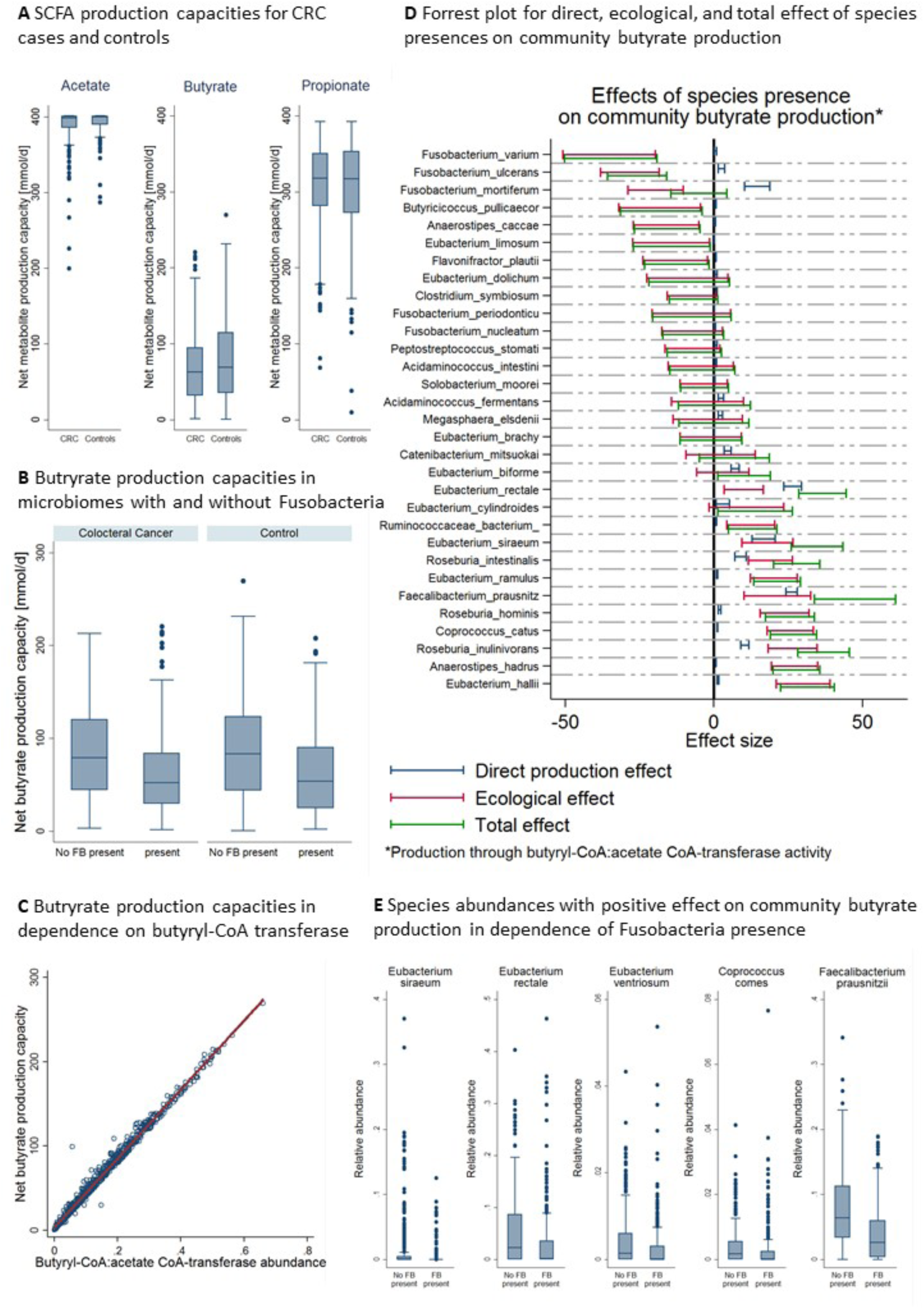
Overview over simulation results regarding short chain fatty acid production. **A** Box plots for acetate, butyrate, and propionate net production capacities for CRC cases and controls. Net production capacities are significantly different across cases and controls for acetate (p=0.046) and butyrate (p=0.030). **B** Box plots for net butyrate production capacities for cases and controls across microbiomes with and without *Fusobacteria* presence. Communities with *Fusobacteria* had significantly lower net butyrate production potentials (p=4.43e-09). **C** Scatter plot with regression line for net butyrate production capacities in dependence on the butyryl-CoA:acetate CoA-transferase abundance (R-squared=0.99). **D** Forest plots for direct, ecological and total effects of species presence on community butyrate production through the butyryl-CoA:acetate CoA-transferase route. Caps represent 95%-confidence intervals. Only species found in at least 5% of all samples were included. **E** Box plots for species abundances positively associated with community butyrate production in dependence of *Fusobacteria* presence. All species were significantly less abundance when Fusobacteria were present in the microbiome (all p<0.001). SCFAs=short chain fatty acids, FB=Fusobacteria. CRC=colorectal cancer.

### *Fusobacteria* species have large negative ecological effect on butyrate production through the butyryl-CoA:acetate CoA-transferase route

To elucidate the changes in the community associated with *Fusobacteria* causal to the lower butyrate production potential, we calculated for each butyrate producing species found in at least 5% and maximally 95% of all samples the direct butyrate production capacity and their ecological effects on the community butyrate production (Methods). Three reactions abundances showed a correlation r>0.99 with the community butyrate production capacity: The conversion reaction of crotonoyl-CoA to butyryl-CoA by Bcd-Etf complex (VMH identifier: BTCOADH), the butyryl-CoA:acetate CoA-transferase (VMH identifier: BTCOAACCOAT), and the ferredoxin:NAD oxidoreductase (VMH identifier: FDNADOX_H). From those three, which belong to the same pathway, the butyryl-CoA:acetate CoA-transferase directly produces butyrate with variance in its abundance being responsible for over 98% of variance in net community butyrate production capacity. Thus, abundance of this reaction directly translates into net butyrate production capacity in a proportional manner (R-Squared=0.99 Fig. 3C), representing thereby the main route for microbial butyrate production in the population of interrogated community models. While all five *Fusobacteria* sp. detected in at least five percent of the samples were predicted to produce small amounts of butyrate via the butyryl-CoA:acetate CoA-transferase route, they had large negative ecological effects on community butyrate production (Fig. 3D, Table S2). *F. varium, F. mortiferum*, and *F. ulcerans* had the highest negative impact on community butyrate production across all modelled butyrate producing species (Fig. 3D). Highlighting the negative impact of *Fusobacteria* presence, from seven species that contributed at least 10% of variance to the net community butyrate production capacity with positive effect sign (Table S3), five were negatively correlated with the presence of *Fusobacteria*, although the effect regarding *Coprococcus comes* missed significance after adjusting for the study group variable (OR=0.70; 95%-CI:(0.48;1.02), p=0.06, Table S4). The effect was most drastic with the well-known fibre degrader *Faecalibacterium prausnitzii* (OR=0.49, 95%-CI:(0.42;0.58), p=8.41e-18, FDR<0.05 Fig. 3E, Table S4), which is known to produce butyrate through the butyryl-CoA:acetate CoA-transferase route (Louis et al., 2014).

### Faecal metabolomics validates community butyrate production predictions

All the results until now are based on *in silico* calculations. Now, we focus on the validation of core results using faecal metabolome data from the same cohort where for 347 individuals, faecal metabolome measurements were available including quantifications for butyrate and glutarate (Yachida et al., 2019). The community models made distinct predictions i) for the net butyrate production capacity, ii) for the species contributing to community butyrate production, and iii) the prediction that butyrate community production is lowered in communities with prevalent fusobacterium species. First, predicted butyrate secretion capacities were significantly correlated with measured log faecal butyrate concentrations (b=0.005, 95%-CI:(0.003,0.006), p=9.87E-10) explaining overall 10.9% of faecal butyrate concentration variance (see Fig. 4A). Second, we calculated the full species butyrate association pattern by regressing the faecal log butyrate concentrations on the species presences in sequential regressions while adjusting for case-control status, age, sex, and BMI. The corresponding *in silico* species-metabolite association statistics were then derived from analogous regressions using the net community butyrate production capacity as response variable. The summary statistics for the species butyrate association patterns *in vivo* and *in silico* can be found in the supplementary material (Table S5). From 47 nominally significant species faecal butyrate associations, community modelling predicted the sign correctly for 43 (prediction accuracy: 91.49%, Fisher’s exact test: p=1.69e-08). From 17 FDR corrected significant species faecal butyrate associations, community modelling predicted in all but one case (*Granuticatella adiacens*) the sign (prediction accuracy: 94.1%, Fisher’s exact test: p=0.006) (Fig. 4B). Beyond the sign, community modelling predictions were additionally significantly correlated with the size of the regression-based association statistics for the nominally significant species (r=0.75, p=9.96e-10) and the FDR corrected significant species (r=0.86, p=7.65e-06) (Fig. 4D). Moreover, as predicted by the modelling, individuals with prevalent *Fusobacterium* species sp. had significantly lower faecal butyrate levels (b= −0.19, 95%-C:(−0.34, −0.05), p=0.011) (see Fig. 4C) despite fusobacteria themselves being butyrate producers, reflecting the predicted deleterious effects of *Fusobacteria* on other butyrate producing species, As the faecal metabolome is considered to be partly a readout of the functional capabilities of the gut microbiome (Yachida et al., 2019; Zierer et al., 2018), this data could provide a proof of principle for the validity of AGORA-based community modelling. However, the variance in the faecal metabolome is also determined by variance in nutrition habits and attributes of the host; both of which were not modelled in this work, thereby limiting the extent to which the variance in the faecal metabolome could be explained by community modelling. Note that the utilised modelling algorithms utilised above were not “trained” in any way on the utilised metabolome dataset. In conclusion, community modelling was able to predict measured species butyrate correlations with high accuracy and, thus, to predict the species-level contribution to the faecal butyrate pool.

**Figure 4:**
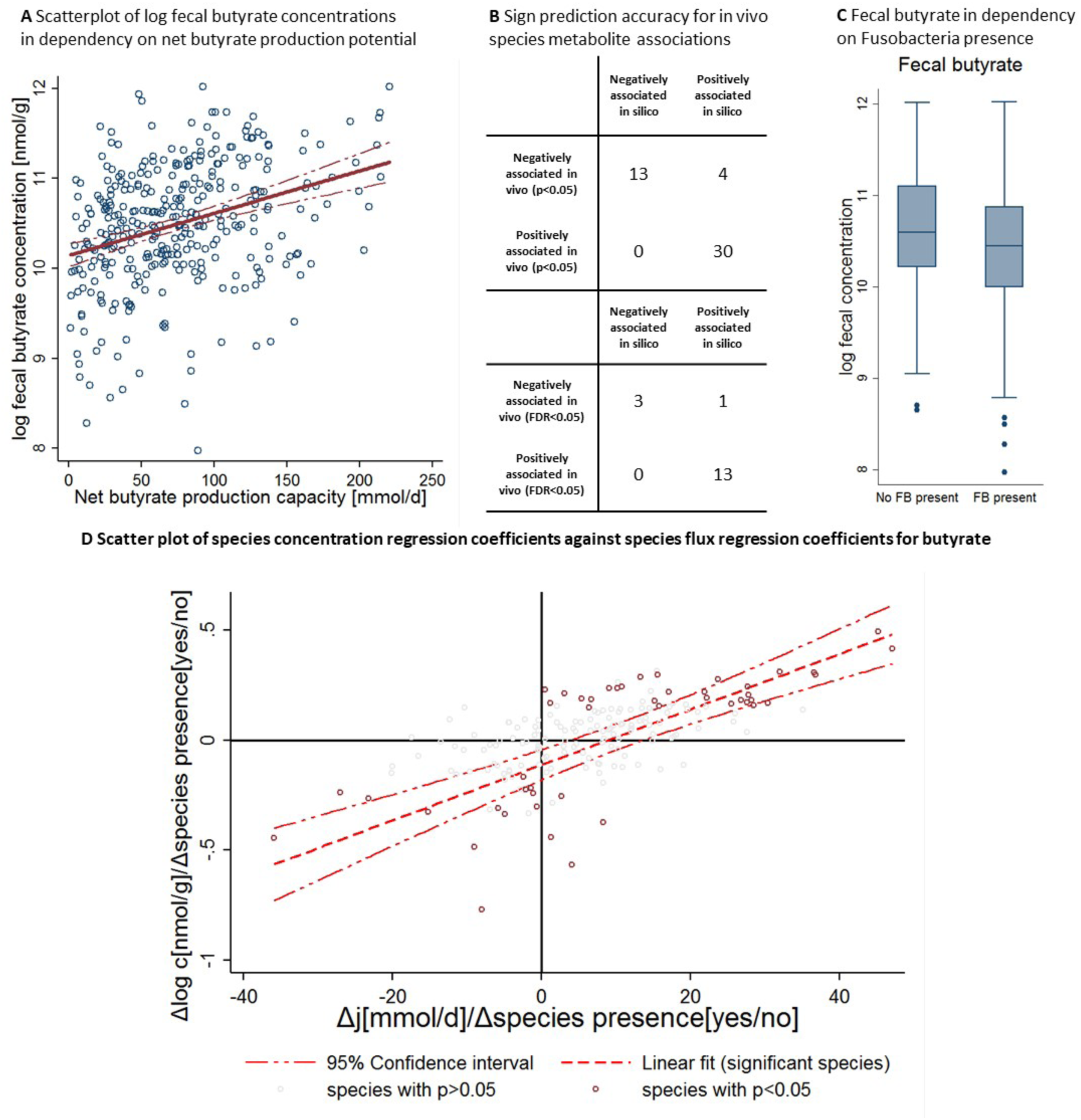
Validation of community modelling predictions regarding butyrate. **A** Scatter plot with regression line of log faecal butyrate concentrations against community net butyrate production capacities. The regression slope is significantly different from zero (b=0.00445, 95%-CI:(.00295,.00595), p=1.22E-08). **B** Accuracy of sign prediction for significant species faecal butyrate concentration association through community modelling. **C** Box plots for log faecal butyrate concentrations for microbiomes with and without *Fusobacteria*. Microbiomes with *Fusobacteria* were associated with lower faecal butyrate levels (b= −0.18, 95%-C:(−0.33, −0.03), p=0.020). **D** Scatter plot with regression line of empirical species faecal butyrate association statistics (expressed as regression coefficients) against *in silico* species net metabolite production association statistics (expressed as regression coefficients). *In silico* and empirical association statistics were significantly correlated with each other (r=0.75, p=9.96e-10). FB=Fusobacteria, c=concentration, j=net production capacity flux.

### Faecal glutarate levels indicate net glutarate consumption by microbial communities

Then, we turned our attention to the relation between *in silico* predicted net glutarate production capacity and actual experimentally measured faecal glutarate concentrations. Surprisingly, we discovered that communities with the capability of glutarate production were associated with significantly lower glutarate levels in faeces (b= −0.44,95%-CI(−0.68, −0.20), p=3.24e-04) (see Fig. 5A), explaining 4.06% of variation in faecal glutarate pools. In consequence, faecal glutarate concentrations were significantly lower in the presence of *Fusobacteria*. Remember that glutarate production capability is synonymous for *Fusobacteria* presence. The microbial transport reaction for glutarate is bi-directional and the necessary reactions of glutarate production co-occur with the degradation reactions leading to butyrate production from glutarate (Fig. 2D). Hence, it is possible that a positive net glutarate production capacity indicates that glutarate can be taken up for ATP generation. In this scenario, communities would be able to consume glutarate, explaining the inverse association of net metabolite production capacity and faecal metabolite concentration. This interpretation is corroborated by testing the ability of community modelling to predict species faecal glutarate associations (Table S6). From 69 nominally significant species faecal glutarate associations, 62 were in line with the community modelling prediction when interpreting the secretion potential as a measure of consumption (prediction accuracy: 89.86%, Fisher’s exact test: p=2.28e-12) (Fig. 5B). For 50 out 56 FDR corrected significant associations, community prediction correctly predicted the sign (prediction accuracy: 89.39%, Fisher’s exact test: p=1.27e-09) (Fig. 5B). As with butyrate, community modelling was also able to predict size of regression coefficient of the species for the faecal glutarate concentration (r=-0.76, p=2.89e-14 for the nominally significant species; r=-0.74, p=5.36e-11 for the FDR corrected species) (Fig. 5D).

**Figure 5:**
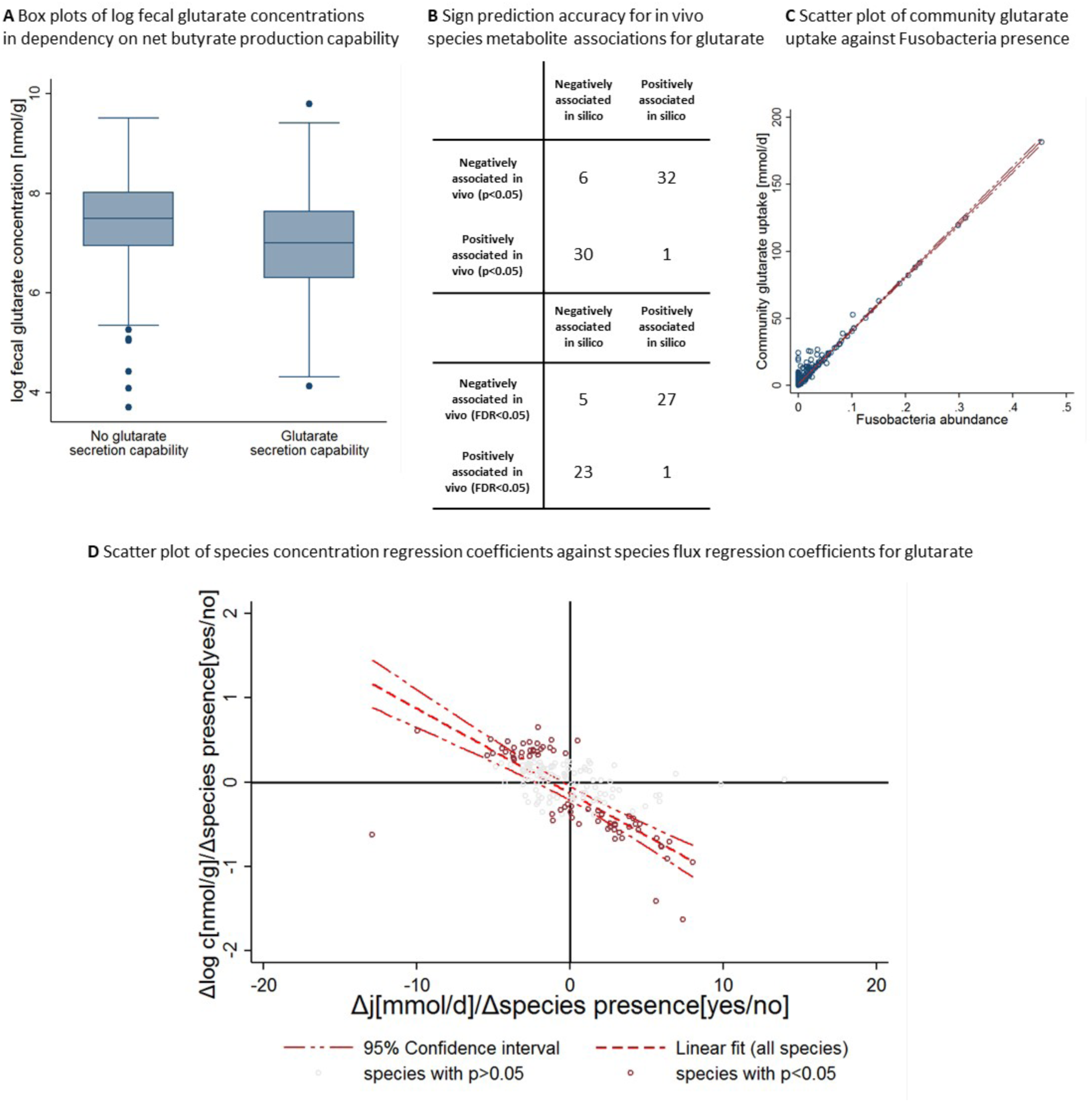
Validation of community modelling predictions regarding glutarate. **A** Box plots for log faecal glutarate concentrations for communities with and without glutarate secretion capability. Communities with glutarate secretion capability are associated with significantly lower faecal glutarate concentrations (b= −0.44,95%-CI(−0.68, −0.20), p=3.24e-04). **B** Accuracy of sign prediction for significant species faecal glutarate concentration association through community modelling. **C** Scatter plot of *in silico* community uptake of glutarate against *Fusobacteria* abundance (r=0.98). **D** Scatter plot with regression line of empirical species faecal glutarate association statistics (expressed as regression coefficients) against *in silico* species net metabolite production association statistics (expressed as regression coefficients). *In silico* and empirical association statistics were significantly correlated with each other (r=-0.76, p=2.89e-14). c=concentration, j=net production capacity flux.

### Faecal glutarate consumption is driven by *Fusobacteria* sp. *in silico*

Above, we showed that community glutarate secretion *in silico* is likely an indicator for glutarate consumption *in vivo*. Testing this interpretation, we designed additional simulations to model the glutarate uptake by those species who are able consume glutarate. Note that while only *Fusobacteria* were able to secrete glutarate, we identified 16 species present in at least one microbiome being able to take up glutarate including the seven detected *Fusobacteria* sp. (Table S7). However, *Fusobacteria* abundance was the primary determinant of glutarate uptake potential (R-squared=0.97, see Fig. 5C). Consequently, uptake potential and community secretion potential for glutarate correlated strongly with each other (r=0.98, p<1e-30, Fig. 5C). In conclusion, the interpretation of the community glutarate production capacity being an indicator of the potential to consume glutarate was also supported by the species level uptake modelling.

## Discussion

A key challenge for a mechanistic understanding of the gut microbiome in health and disease is to map Changes in gut microbial abundances onto functional changes impacting the host’s metabolism. Here, we present a functional metabolic modelling approach combining COBRA modelling with population statistics that enables translating individual-specific microbial abundances into personalised microbial metabolite profiles. Through this framework, we demonstrated that each person’s gut microbiome is functionally unique, emphasising the need for individualised modelling of microbiomes as possible with COBRA community modelling. We highlighted the utility of our approach by generating insights on the functional alterations associated with *Fusobacteria* sp. presence in the gut microbiome; insights of potential clinical relevance especially in CRC where *Fusobacteria* sp. are enriched (Kostic et al., 2013; Mehta et al., 2017; Zhou et al., 2018). Finally, we validated the prediction of the *in silico* modelling against faecal metabolome data, revealing excellent agreement between *in silico* predictions and empirical data.

Our analyses of net production capacities revealed alterations in the domain of fermentation products in CRC, including short chain fatty acids. CRC microbial communities had lower net production capacities in acetate and butyrate (Fig. 3). The lower production capacity of short chain fatty acids is of potential clinical relevance due to the known anti-inflammatory, anti-tumour-effects of butyrate (Koh et al., 2016). Moreover, butyrate is a main energy source for colonocytes but not for cancer cells, which prefer glucose (Koh et al., 2016). Evidence exists for butyrate having protective properties for colon-cells and low fibre intake has been considered as a risk factor for CRC (Chang et al., 2014). The finding that CRC microbiomes have decreased capacities in producing butyrate fits with earlier observations of depletion of butyrate producing species in CRC microbiomes (Wu et al., 2013; Zhu et al., 2014).

While well documented, the cause for the depletion of butyrate producing species in CRC is less understood. In our study, we found that presence of *Fusobacteria* sp. is strongly associated with this shift in the community composition, quantified by the high negative ecological effect of *Fusobactera* sp. on community butyrate production (Fig. 3). Importantly, the negative effect of *Fusobactera* sp. is not a CRC specific feature: In healthy individuals, the presence of *Fusobacteria* was associated with lower butyrate production capacities as well (Fig. 3B). This observation fits well with *in vitro* studies showing that *F. nucleatum* produces bactericidal compounds hazardous to butyrate-producing species, in this case *F. prausnitzii* (Guo et al., 2018). It should be noted that the highest negative effects on community butyrate production were with *F. varium, F. mortiferum*, and *F. ulcerans*, indicating that not only *F. nucleatum* may play a role in CRC (Fig 3D). Noteworthy, *Fusobacteria* sp. co-occur with each other (Zhou et al., 2018), making inferences on single species complicated. For example, in the present study, we also found *F. mortiferum* to be significantly enriched in CRC (Fig. 2C). In conclusion, the evidence points overall towards *Fusobactera* sp. being deleterious for community butyrate production.

*F. nucleatum* has been repeatedly implicated in CRC (Flanagan et al., 2014; Mima et al., 2016; Ng et al., 2019). While it has been described that *F. nucleatum* plays a role in treatment resistance in CRC and in the modulation of anti-tumour inflammation response (Mima et al., 2015; Yu et al., 2017), the metabolic role of an enrichment in *F. nucleatum* and in other *Fusobacteria* sp. in CRC is less clear. In this respect, we found a clear enrichment of the capability to produce glutarate from lysine in CRC microbiomes, which is mechanistically linked to *Fusobacteria* presence (Fig. 2D). Importantly, this feature is a metabolic trait of all seven *Fusobacteria* that we detected in this study, and a general feature of all species in the Fusobacteria genus captured in the VMH resource (Noronha et al., 2019). *Fusobacteria sp*. are the only species in the AGORA collection having the full pathway from lysine to glutarate *and* an exchange reaction for glutarate. In line with our study, Fusobacteria are known for their asaccharolytic metabolism (Flynn et al., 2016). As glutarate is an intermediate on the pathways from lysine and from glutarate to butyrate (Vital et al., 2014), this suggests that the increased *Fusobacterium* abundance in CRC microbiomes would result in increased amino acid fermentation, in particular lysine to butyrate. An enrichment in amino acid degradation pathways accompanied by a corresponding decrease in carbohydrate degradation has been reported for CRC microbiomes (Wirbel et al., 2019), fitting with our results. It is noteworthy that we found *Fusobacteria* species to be enriched in men. Men have higher risks for developing CRC (White et al., 2018), sparking the speculation whether Fusobacteria presence may mediate a part of the sex-specific risk for CRC, although the discussion around sex-differences in CRC are complicated by social and cultural effects (Kim et al., 2015).

Interestingly, integration with faecal metabolomics indicated that *Fusobacteria* are likely net consumer of glutarate and the main determinant of community glutarate uptake. Glutarate, however, is biochemically closely related to alpha-ketoglutarate and thereby to the Krebs cycle. Aberrations in the Krebs cycle in return are a hallmark of cancer metabolism (Anderson et al., 2018; Pavlova and Thompson, 2016). Thus, CRC metabolism may be interlinked with the metabolism of *Fusobacteria*, allowing for the speculation that *Fusobacteria* may profit from Krebs cycle alterations in CRC.

Previously, we have demonstrated the use of personalised metabolic modelling for the stratification of paediatric inflammatory bowel disease patients and controls in a *purely in silico* approach (Heinken et al., 2019) and validated changes in the metabolome of Parkinson’s Disease patients with personalised models built from an unrelated cohort (Hertel et al., 2019). Here, by integrating the AGORA based COBRA community modelling predictions with faecal metabolomics, we could validate our predictions regarding butyrate, glutarate, *Fusobacteria* and other butyrate producing species. We were able to correctly predict, which species correlate with faecal butyrate and glutarate levels, and even the effect sizes of these associations were predicted correctly to a high degree (Fig. 4, 5). This functional metabolic modelling delivers a new proof of principle for community modelling, opening new routes of applications. As butyrate production is considered to be integral for gastrointestinal health (Chang et al., 2014), probiotic, prebiotic, and synbiotic interventions have started targeting beneficial butyrate producers, such as *F. prausnitzii* (Chang et al., 2019). AGORA-based community modelling enables the prediction of the outcome of therapeutic and dietary interventions (Thiele et al., 2017). Our study now reveals that these *in silic*o biomarkers are indeed reflective of the gut microbiome’s metabolic capacities and in good agreement with faecal butyrate concentrations. Importantly, the models were not contextualised with the metabolome data from Yachida et al. during their construction, meaning that the Yachida et al. dataset delivers an external validation (Yachida et al., 2019). Thus, *in silico* modelling can deliver computational biomarkers for phenotypes, which could be used, in principle, for diagnostic or prognostic purposes. Additionally, our work highlights that community modelling can be utilised as a further layer of validation for empirical species metabolome association studies where correlations are often difficult to interpret due to uncontrolled confounding (Noecker et al., 2019). As community modelling is based on deterministic calculations from microbiome measurements, certain types of confounding have no effect on *in silico* species metabolite association. Thereby, community modelling can help in diminishing false positives in microbiome metabolome association studies; an important aspect as noted in earlier work (Noecker et al., 2019).

While the modelling was overall in good agreement with the empirical metabolome measurement, several limitations should be noted. We applied one standardised diet, excluding therefore variance caused by differential diet habits from the analyses. However, the general methodology would allow the personalisation of the diet information used for modelling. Thus, if the diet habits are sampled in a suitable way, the type of calculation performed here can be individualised not only regarding microbial abundances, but also regarding the diet information (Baldini et al., 2018). Furthermore, this study did not integrate the host’s metabolism into the modelling. Further studies, based on whole body organ resolved COBRA modelling (Thiele et al., 2020), could deliver more insight into the interplay between the host and the microbiome in CRC and beyond. Knowledge about microbial functions and genomic annotations are incomplete, and as such, the AGORA collection is subject to constant updates. Another known limitation of COBRA is the lack of kinetic parameters and the simulation of fluxes rather than concentrations due to the steady-state assumption. However, the good agreement between *in silico* fluxes and experimentally measured concentrations in this study suggests that is it possible to mechanistically translate increased or decreased fluxes into increased or decreased concentrations. Importantly, this study is based on cross-sectional data and as such, causality between clinical parameters and microbial functions cannot be established. However, determinations of metabolic functions by community modelling are not confounded by factors like age, sex, exercise or other factors, as they are deterministic calculations from abundance patterns. Providing a major conceptual advantage regarding the functional analyses of species metabolite associations via calculating abundance concentrations correlations, community modelling allows for the dissection of direct contributions of species to and their ecological effects on the community metabolite production capacities. Noteworthy, ecological effects, as defined in this work, allow the mapping of the statistical effects of the presence of species on the community structure in terms of metabolic function. It should, however, be remembered that ecological effects note statistical associations and are not necessarily of causal nature.

In conclusion, AGORA-based community modelling provides a powerful toolset for the characterisation of microbial metabolic functions in health and disease, delivering testable hypotheses, *in silico* biomarkers, and potential endpoints for clinical studies. Importantly, the AGORA reconstructions had been extensively curated based against comparative genomics and experimental data two microbial textbooks and over 200 peer-reviewed papers (Magnusdottir et al., 2017). Thus, underneath the conclusions presented in this paper lies accurate, manually gathered knowledge on fermentation pathways in hundreds of organisms. Overall, this study provides a proof of principle that the knowledge encoded in the AGORA models can be translated into clinical insight via community modelling.

## Author contributions

J.H., A.H, and I.T. designed the study. J.H., A.H. wrote the manuscript. J.H. performed the statistical analyses and developed the theoretical concepts. A.H. performed the community modelling. A.H. and F.M. designed the community models. All authors reviewed and approved the final manuscript.

### Acknowledgements

The study was funded by the European Research Council (ERC) under the European Union’s Horizon 2020 research and innovation program (grant agreement No 757922).

## Methods

### Study sample

We utilised the Japanese colorectal cancer cohort data from (Yachida et al., 2019), which had publicly available shotgun sequencing data for n=616 individuals (365 CRC cases and 251 healthy controls). The reads had already been processed and taxonomic profiling utilizing MetaPhlAn2 (Truong et al., 2015). Attached to this dataset several, meta-data on age, sex, BMI, smoking, alcohol, stages of the disease, and tumour location were available. Additionally, linked to these data, faecal metabolome quantifications were available for n=347 probands (CRC: 220, controls:127), allowing the validation of attributes of the community models by linking them to empirical metabolome quantifications. For details on metagenomic and metabolomic measurements, refer to (Yachida et al., 2019).

### Definition of an average Japanese diet

An average Japanese diet was defined based on the mean daily food consumption in 106 Japanese extracted from food frequency questionnaires and 28 days weighed diet records (Tokudome et al., 2001) (Table S8a). Therefore, we used the Diet Designer of the VMH database (https://vmh.life), which lists the composition of >8,000 food items (Noronha et al., 2019). In the absence of a perfect match, the most related food item entries were retrieved. The Diet Designer calculates uptake flux values in mmol/person/day for each nutrient component based on the specified diet, as described elsewhere (Noronha et al., 2019). We integrated these uptake flux values as diet constraints with all community microbiome models using the Microbiome Modelling Toolbox (Baldini et al., 2018) (see below). To ensure that all AGORA pan-species models could grow under the defined diet, we adapted the calculated uptake fluxes as necessary (Table S8b). The diet constraints were defined to be in mmol/person/day.

### Simulations

All simulations were performed in MATLAB (Mathworks, Inc.) version R2018b with IBM CPLEX (IBM) as the linear and quadratic programming solver. The simulations relied on functions implemented in the COBRA Toolbox (Heirendt et al., 2019), and the Microbiome Modelling Toolbox (Baldini et al., 2018).

### Construction of sample-specific gut microbiota models

Metagenomic datasets from 616 samples were used as published in (Yachida et al., 2019). We utilised the sequencing data from the corresponding supplementary material (https://static-content.springer.com/esm/art%3A10.1038%2Fs41591-019-0458-7/MediaObjects/41591_2019_458_MOESM3_ESM.xlsx). The data had been already preprocessed and available in relative abundances on the species level. The relative abundances were then mapped onto the reference set of 773 AGORA genomes (Magnusdottir et al., 2017) through the translateMetagenomeToAGORA.m function in the Microbiome Modelling Toolbox (Baldini et al., 2018). Via the mgPipe module of the Microbiome Modelling Toolbox, personalised microbiome models were derived. In brief, the corresponding AGORA reconstructions of all strains found in at least one microbiome were put together into one global constraint-based microbiome community reconstruction as described before (Baldini et al., 2018; Thiele et al., 2013). Then, the biomass objective function was coupled with the flux through each AGORA species panmodel (for details see (Heinken et al., 2013)), parametrising the community biomass reaction via the relative abundances as stoichiometric values for each microbe biomass reaction in the community biomass reaction. The models were appropriately contextualised with the average Japanese Diet described above. The resulting diet exchange fluxes were then applied to community models (Baldini et al., 2018). The flux through the community biomass reaction was set to be between 0.4 and 1 mmol/person/day, as described before. The features of the personalised community models are given in Fig. 1.

### Definitions and theoretical frameworks

#### Utilised attributes of populations of community models

Let be *M* = {*M*_1_, *M*_2_, … *M*_*I*_} a set of *I* community models corresponding to *I* measured microbiomes. We are interested here in three attributes of the model *M*_*i*_:

i. the vector of microbial abundances ***a***_***i***_ ∈ [0,1]^*K*^ belonging to the model *M*_*i*_ where *K* denotes the number of species included into the AGORA collection.
ii. the vector of reaction abundances ***r***_***i***_ ∈ [0,1]^*J*^ belonging to the model *M*_*i*_ where *J* denotes the number of reactions included into the AGORA collection in total.
iii. A vector of net metabolite production capacities ***n***_***i***_ ∈ [0, *c*_*l*_]^*L*^ with *c*_*l*_ being the maximum possible net metabolite production capacity under the set of applied constraints and L being the number of metabolites with microbial exchange reactions in at least one AGORA genome scale model. Net metabolite production capacities are defined by the difference of maximal secretion and maximal uptake fluxes. We say that a model *M*_*i*_ has a net production capability for the metabolite *l* if *n*_*il*_ > 0.

Thus, our population statistics analyses of community models were performed on microbial abundances, reaction abundances, net metabolite production capacities and net metabolite production capabilities.

#### Metabolic Equivalence

Now, we define the term metabolic equivalence, which allows us to cluster microbial communities having the same set of metabolic functions.

##### Definition 1: Metabolic Equivalence

We call two community models *M*_*j*_ and *M*_*k*_ metabolic equivalent regarding the (sub)set *E* of metabolites with exchange reactions in at least one AGORA genome scale model if and only if that for all *l* ∈ *E* it holds that *n*_*jl*_ > 0 ⇔ *n*_*kl*_ > 0. We then write *M*_*j*_ ∼_*E*_ *M*_*k*_.

This defines an equivalence relation, as the relation ∼_*E*_ fulfills the attributes of being reflexive (*M*_*j*_ ∼_*E*_ *M*_*j*_), symmetric (*M*_*j*_ ∼_*E*_ *M*_*k*_ ⇔ *M*_*k*_ ∼_*E*_ *M*_*j*_), and transitive (*M*_*j*_ ∼_*E*_ *M*_*k*_ *and M*_*k*_ ∼_*E*_ *M*_*l*_ ⇒ *M*_*j*_ ∼_*E*_ *M*_*l*_).

#### Necessary and sufficient conditions for net metabolite production capacities

Now, we define *sufficient* and *necessary* attributes for net metabolite production capabilities given a set of microbial community models *M* = {*M*_1_, *M*_2_, … *M*_*I*_}. The concepts of *“metabolically sufficient”* and *“metabolically necessary”* will be analogous for species and reactions. First, however, we will define *informative metabolites*.

##### Definition 2: Informative metabolite

We call a metabolite *l* informative, if and only if ∃*M*_*i*_ ∈ *M*: *n*_*il*_ > 0 *and* ∃*M*_*j*_ ∈ *M*: *n*_*jl*_ = 0. Informative metabolites are therefore those metabolites with variance in the net production capabilities across the set of models *M*.

##### Definition 3: Necessary and sufficient reactions

Let be *l* an informative metabolite. Then, we call a reaction *k* necessary if and only if for all *M*_*i*_ ∈ *M* it holds that *r*_*ik*_ = 0 ⇒ *n*_*il*_ = 0. We call a reaction *k* sufficient if and only if for all *M*_*i*_ ∈ *M* it holds that *r*_*ik*_ > 0 ⇒ *n*_*il*_ > 0.

##### Definition 4: Necessary and sufficient species

Let be *l* an informative metabolite. Then, we call a species *j* necessary if and only if for all *M*_*i*_ ∈ *M* it holds that *a*_*ij*_ = 0 ⇒ *n*_*il*_ = 0. We call a species *j* sufficient if and only if for all *M*_*i*_ ∈ *M* it holds that *a*_*ij*_ > 0 ⇒ *n*_*il*_ > 0.

Thus, we call species and reactions *necessary* for a certain metabolic function, if their absence implies missing the metabolic function under consideration in all observed community models. In contrast, we call species and reactions *sufficient* for a metabolic function, if their presence implies showing the metabolic function of interest in all models. It is important to note that the concepts of necessity and sufficiency are defined for metabolites, which are neither produced by all models, nor by any of the models. We can only learn *necessary* and *sufficient* conditions from variance in the occurrence, which motivates the definition of *informative metabolites*. This is in parallel to statistics where variance in the random variables is a prerequisite to identify patterns of stochastic dependency. As in statistics, the dependency relations given by sufficiency and necessity should not be confused with causality, as conditions could co-occur in the set communities observed. Therefore, we define the concepts of strictly sufficient and strictly necessary, which introduces a type of conditional dependence notion.

##### Definition 5: Strictly necessary reactions

Let be *l* an informative metabolite. Let be *Q*_*l*_ the set of all reactions, which are necessary for net production capability for the metabolite *l* and *k* ∈ *Q*_*l*_ a specific necessary reaction. We call *k* strictly necessary if and only if ∃*M*_*i*_ ∈ *M* with *r*_*ik*_ = 0 and ∀j ∈ *Q*_*l*_\*k*: *r*_*ij*_ ≠ 0.

##### Definition 6: Strictly necessary species

Let be *l* an informative metabolite. Let be *Q*_*l*_ the set of all species, which are necessary for net production capability for *l* and *k* ∈ *Q*_*l*_ a specific necessary species. We call *k* strictly necessary if and only if ∃*M*_*i*_ ∈ *M* with *a*_*ik*_ = 0 and ∀j ∈ *Q*_*l*_\*k*: *a*_*ij*_ ≠ 0.

##### Definition 7: Strictly sufficient reactions

Let be *l* an informative metabolite. Let be *Q*_*l*_ the set of all reactions, which are sufficient for net production capability for *l* and *k* ∈ *Q*_*l*_ a specific sufficient reaction. We call *k* strictly sufficient if and only if ∃*M*_*i*_ ∈ *M* with *r*_*ik*_ > 0 and ∀j ∈ *Q*_*l*_\*k*: *r*_*ij*_ = 0.

##### Definition 8: Strictly sufficient species

Let be *l* an informative metabolite. Let be *Q*_*l*_ the set of all species, which are necessary for net production capability for *l* and *k* ∈ *Q*_*l*_ a specific necessary species. We call *k* strictly necessary if and only if ∃*M*_*i*_ ∈ with *a*_*ik*_ > 0 and ∀j ∈ *Q*_*l*_\*k*: *a*_*ij*_ = 0.

It is important to realize that the definitions presented here are dependent on the variance in the population of microbial communities. The larger the sample size, the more necessary and sufficient conditions will be discovered. Sufficiency and necessity are technical attributes of populations of community models in the first place. The identified conditions do not need to be necessary and sufficient in a biological sense. However, they are valuable candidates for being indicators of causal processes and thus targets for experimental validation.

#### Direct, ecological, and total effect of species on net community metabolite production capacities

Here, we define formally the effects of a presence of a species on the net community metabolite production capacities observed in a population of community models *M*. The concepts of effects are defined via populations statistics. Therefore, these concepts must be treated as statistical estimates and should always be reported with confidence intervals.

##### Definition 9: Average direct species net production effect

Let l be a metabolite and M the population of community models. The average direct species production effect 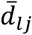 for a metabolite l and a species j is defined by

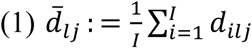

where *d*_*ilj*_ stands for the net production (through secretion and uptake) of the metabolite l by the species j in the community model *M*_*i*_. We call *d*_*ilj*_ the species net production capacity.

A species, however, cannot only impact the net community production capacity by direct contributions. A species can also impact the production of other microbes and can be associated with alteration in the community structure, changing the abundance of other microbes relevant for the community production of a metabolite. This motivates the definition of the ecological species effect, which gives a measure of these indirect influences associated with the presence of a microbe.

##### Definition 10: Ecological species effect

Let *l* be a metabolite and *M* the population of community models. Let *M*^*j*^ ≔ {*M*_*i*_: *a*_*ij*_ > 0} be the set of community models with the abundance of the species *j* greater than zero, and *M*^¬*j*^ ≔ {*M*_*i*_: *a*_*ij*_ = 0} the set of community models missing the species *j*. The ecological species effect *ē*_*lj*_ is then given by

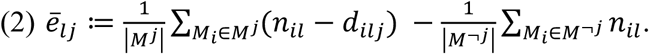

Thus, the ecological species effect is the difference between average net metabolite production capacities of communities with a species and communities without a species after discounting the direct species net production capacity. Note that the direct species net production is zero in all models belonging to the set *M*^¬*j*^.

Obviously, the ecological species effect is not necessarily causal, and it can be calculated conditional on a set of covariates minimising confounding by basic covariates, such as age, sex, or BMI, via multivariable regressions.

##### Definition 11: Total species effect

Let *l* be a metabolite and *M* the population of community models. Then, the total species effect 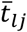 is defined by the sum of average direct species net production effect and the ecological species effect:

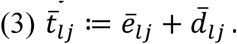

The total species effect is the difference in net production capacities between the community models having a certain species and the community models missing this specific species.

### Statistical analyses

We performed statistical analyses on the following attributes of community models: 1) net metabolite production capabilities, 2) net metabolite production capacities, 3) reaction abundances, and 4) species abundances. Due to one infeasible model, the final sample size for analysing relations between metadata and attributes of the community models was n=615 and the final sample size for analysing the community models together with the faecal metabolome was n=346. For descriptive statistics, metric variables were expressed in means and standard deviations, categorical variables were described by proportions. All p-values are reported two-tailed. The statistical analyses were performed with STATA 14/MP (STATA Inc., College Station, Texas, USA).

#### Analyses of net metabolite production capabilities

To investigate the potential differences in net metabolite production capabilities between cases and controls, we fitted logistic regressions with the net metabolite production capability as binary response variable (can be produced vs. cannot be produced). The predictor of interest in these logistic regressions was the group variable (binary: CRC cases vs. controls) and age, sex, and BMI were used as covariates to minimize confounding. We analysed only metabolites for which at least 5% and maximally 95% of all community models could produce those metabolites to avoid unstable statistical estimates due to low case numbers. Forty-four metabolites fulfilled this criterion. Accordingly, we corrected for multiple testing using the false discovery rate (FDR) (Benjamini, 2010), acknowledging 44 significant tests. An FDR of 0.05 was chosen as significance threshold.

In a second series of logistic regressions, we checked for associations of net metabolite production capabilities with the CRC stage. Thus, we performed logistic regressions as before exchanging the study group variable for the stage variable (categorical: surgery, multiple polyps, stage 0, stage I/II, stage III/IV) excluding healthy controls from the analysis. The stage variable was then tested on significance using a standard Wald test (Harrell, 2001). Once again, we corrected for multiple testing using the FDR, adjusting the significance threshold for 44 tests. Summary statistics for both series of logistic regressions can be found in supplementary Table S1.

Post hoc, glutarate production capability, being the main result of the screening described above, was checked on associations with basic covariates. To check for association with age and sex, a logistic regression with the net glutarate production capability as response variable was fitted using age and sex as predictors of interest, while adjusting for the study group variable (binary: CRC cases vs. controls). To check for association with BMI, a logistic regression with the net glutarate production capability as response variable was fitted using the BMI as predictor of interest, while adjusting for, age, sex, and the study group variable (binary: CRC cases vs. controls).

#### Analyses of net SFCA production capacities

Next, we tested the association of CRC with net production capacities of SFCAs, namely acetate, butyrate, and propionate. To this end, we fitted linear regressions using the respective net SFCA production capacity as response variable, the study group variable (binary: CRC cases vs. controls) as predictor of interest, and age, sex, and BMI as covariates. Heteroscedastic standard errors were applied in the main analyses. For sensitivity analysis, non-parametric bootstrap-derived confidence intervals were calculated using 2000 replications, but the results remained virtually unchanged. Next, we tested net SFCA production capacities on association with the presence of *Fusobacteria* species. Once again, we used linear regressions as before using this time the presence of Fusobacteria (binary: *Fusobacteria* present vs. Fusobacteria not present) as predictor of interest, correcting for age, sex, BMI and study group by including them as covariates. Additionally, we ran mediation according to the Sobel-Goodman test (PMID: 18697684) testing whether Fusobacteria presence mediated the effect of CRC on net butyrate production capacities. Confidence intervals for the indirect and direct effects were calculated by bootstrapping using 2000 replications.

#### Analyses of direct species production effects and ecological species production effects regarding butyrate

To calculate direct and ecological species effects regarding butyrate, we first screened all reaction abundances on correlation with the net community butyrate production capacities, finding the butyryl-CoA:acetate CoA-transferase as one of the top hits. Then, we derived for 31 species (found in at least 5% and maximally 95% of all samples), the direct species production effect, the ecological species effect, and the total species effect on net community butyrate production through the butyryl-CoA:acetate CoA-transferase. The direct species production effect was calculated by using the regression equation of the butyryl-CoA:acetate CoA-transferase net community butyrate production relation replacing the butyryl-CoA:acetate CoA-transferase abundance by the species abundance. This is justified as butyryl-CoA:acetate CoA-transferase abundance is the sum of all species abundances having the butyryl-CoA:acetate CoA-transferase reaction. Then, the ecological species and the total species effects for the 31 species were calculated according to the equations (2) and (3). Finally, 95%-CIs were calculated for all effects using standard procedures for estimating CIs for arithmetic means. The results then were visualised by a forest plot.

To illustrate the effects of Fusobacteria sp., we explored the effect of Fusobacteria presence on those species, which had the highest positive effect on community butyrate production, contributing at least 10% of variance with positive effect sign. Seven species (*Copprococcus comes, Eubacterium rectale, Eubacterium siraeum, Eubacterium ventriosum, Faecalibacterium prausnitzii, Roseburia intestinales, and Roseburia inulinivorans*) fulfilled these criteria. Then, we fitted a series of seven fractional logistic regressions (Baldini et al., 2020) with the abundance of the seven species as response variables, the presence of *Fusobacteria* sp. (binary: present vs. not present) as predictor of interest, while adjusting for age, sex, BMI, and study group. We corrected the significance level for multiple testing using the FDR, adjusting the significance levels for seven tests. Full results and summary statistics can be found in the supplementary material (Tables S2-S4).

#### Statistical integration of community modelling with faecal metabolomics

To validate the simulation results regarding glutarate and butyrate, we integrated the simulation data systematically with faecal metabolome measurements in 347 individuals of the same cohort, including quantifications of glutarate and butyrate concentrations (Yachida et al., 2019). Note that the faecal metabolome is a representative of human metabolism, diet intake, and microbial metabolism such that it cannot be expected that the microbiome can fully explain variegation in faecal metabolite profiles. However, as the microbiome is one source of variance in faecal metabolite content and the simulations predict systematic variance in metabolite output of the microbiome across individuals, we expect that the association pattern between microbes and metabolite production capacities is reflective of the association pattern between microbes and faecal metabolite concentrations. For statistical analyses, faecal glutarate and butyrate concentrations were log-transformed, minimising the skewness of the distributions.

First, we regressed the measured faecal butyrate and glutarate concentrations on the net community production capacities via linear regressions, including age, sex, BMI, and the study group variable as covariates. In the case of glutarate, we also included the net production capability (binary: can be produced vs. cannot be produced) into the regression model, as only 52% of all models had a net production capacity bigger zero. We evaluated then the predictive value of the net community production capacity, respectively, capability by testing their regression coefficients on zero and calculating the incremental R-squared values (increase in model fit by adding net production capacity/capability variables).

Next, we calculated the full species faecal butyrate concentration association pattern by running linear regressions with the measured faecal butyrate concentration as response variable, the species presence (binary: species present vs. species not present) as predictor of interest, while including age, sex, BMI, and the study group variable as covariates. Heteroscedastic standard errors were used. These regressions were run for all species, which were detected in at least 5% and maximally 95% of all samples, resulting in 181 regressions. We retrieved the regression coefficient of the species presence, the corresponding p-value, and the FDR correcting for 181 tests. In a second step, we derived in the same way the full species net community butyrate production capacity association pattern. Note that the *in silico* association pattern was derived on the full sample n=615, assuming implicitly that faecal metabolome measurements were missing completely at random. Then, we checked for all species faecal butyrate concentration associations with p<0.05, respectively, FDR<0.05, whether the sign of the *in silico* derived regression coefficient for the species butyrate association predicted the sign of the empirically derived regression coefficient via Fisher’s exact test. Moreover, we correlated the two species-butyrate association statistics with each other and tested the Pearson correlation via the standard test on significance. A significant prediction of sign and size of empirically derived regression coefficients was interpreted as a validation of the community modelling. We repeated the same methodology for glutarate.

Summary statistics for the full glutarate and butyrate association patterns, *in silico* as well as *in vivo*, can be found in the supplementary material (Table S5, S6).

## Supplemental figure for the manuscript

**Figure S1:**
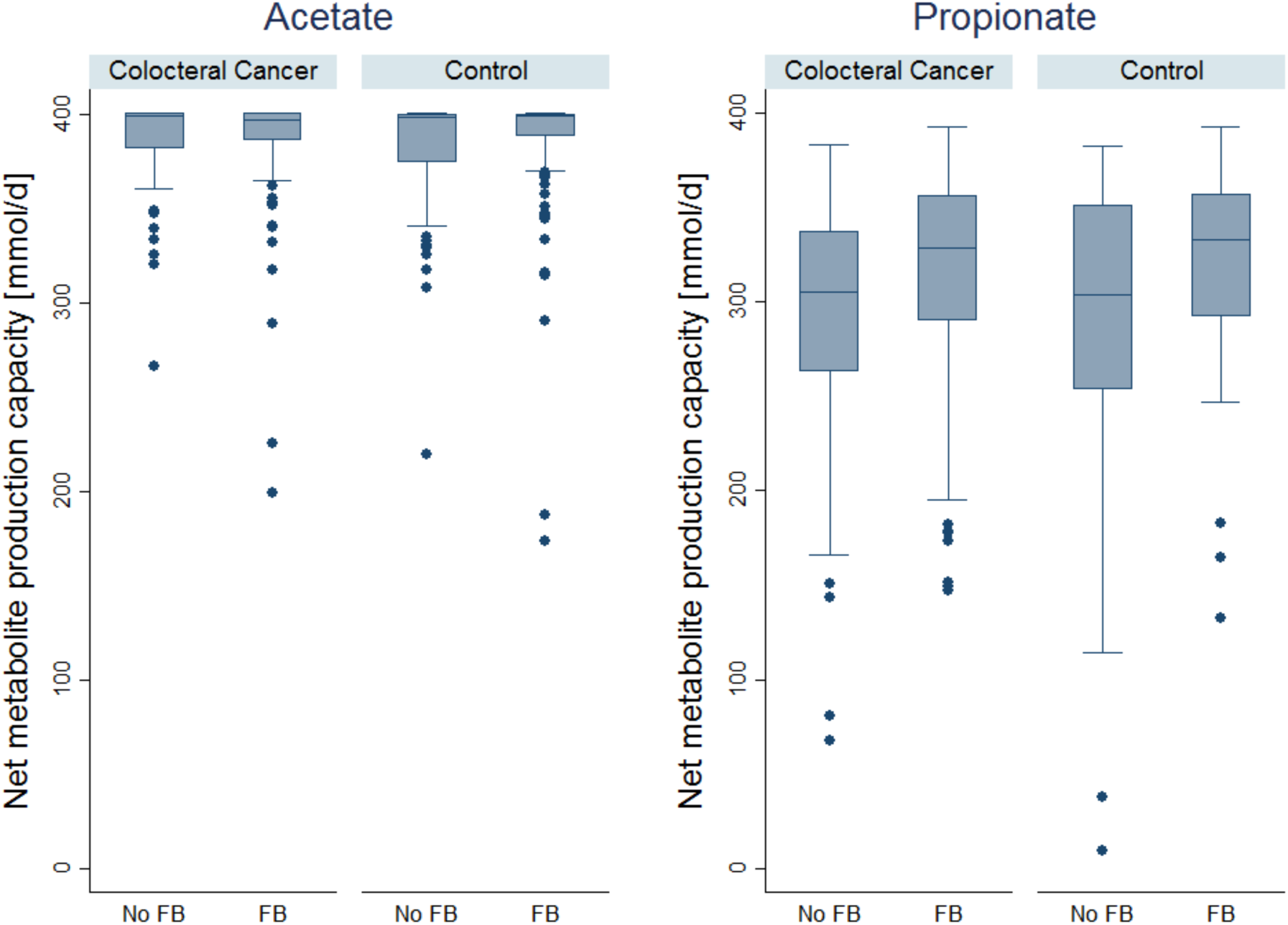
Box plots for net acetate and propionate production capacities for cases and controls across microbiomes with and without *Fusobacteria* presence. Communities with Fusobacteria had significantly higher net propionate production capacities (p=4.87e-07), while acetate production capacities were not significantly different (p=0.519). FB=Fusobacteria.

